# Liver inflammation before spinal cord injury worsens neuropathology, hepatic injury, metabolic syndrome and locomotor deficits

**DOI:** 10.1101/2020.07.15.204578

**Authors:** Matthew T. Goodus, Kaitlin E. Carson, Andrew D. Sauerbeck, Priyankar Dey, Anthony N. Alfredo, Phillip G. Popovich, Richard S. Bruno, Dana M. McTigue

**Affiliations:** The Belford Center for Spinal Cord Injury, Thapar Institute of Engineering and Technology, Patiala, Punjab, India; Department of Neuroscience, Wexner Medical Center, Ohio State University, Thapar Institute of Engineering and Technology, Patiala, Punjab, India; Department of Neurology, Washington University in St. Louis, Thapar Institute of Engineering and Technology, Patiala, Punjab, India; Department of Biotechnology, Thapar Institute of Engineering and Technology, Patiala, Punjab, India; Department of Human Sciences, College of Education and Human Ecology, The Ohio State University, Columbus, OH, USA

## Abstract

Liver inflammation can enhance acute leukocyte recruitment to sites of central nervous system (CNS) injury. The consequences of hepatic inflammation on recovery after injury, however, are unknown. Here, we hypothesize that liver inflammation at the time of spinal cord injury (SCI) will exacerbate spinal cord pathology and impair recovery. Rats receiving SCI with concomitant liver inflammation had worse intraspinal pathology and greater locomotor deficits than rats with baseline liver inflammation. Hepatic inflammation also potentiated SCI-induced non-alcoholic steatohepatitis (NASH), endotoxemia, insulin resistance and adiposity. Circulating and cerebrospinal levels of Fetuin-A were higher in SCI rats with liver inflammation. When microinjected into intact spinal cords, Fetuin-A caused robust macrophage activation, iron accumulation and neuron loss. These studies verify that outcomes from CNS injury are worse if the liver is concomitantly inflamed and implicate Fetuin-A as a potential neuropathological mediator. These novel data suggest hepatic inflammation is a potential clinical target for improving recovery from SCI.

## INTRODUCTION

Major trauma in the body prompts the liver to initiate an acute phase response (APR) that mobilizes inflammatory cells to the injury site. Surprising, however, is that leukocytes infiltrate the *liver* before others enter the injured tissue. For instance, after spinal cord injury (SCI), monocytes and neutrophils invade the liver within 0.5-2 h compared with infiltration of the injured spinal cord by 6 h – 3 d post-injury (Popovich *et al*., 2001; Kigerl, McGaughy and Popovich, 2006; Campbell *et al*., 2008; Anthony and Couch, 2014). This phenomenon in SCI is analogous to the hepatic inflammatory response that exacerbates injury-induced inflammation of the brain. For example, hepatic-specific inhibition of NF-κB signaling after focal brain injury reduced neutrophil infiltration into the brain whereas potentiating hepatic NF-κB signaling enhanced it (Campbell *et al*., 2008). Depletion of hepatic macrophages (Kupffer cells) prior to SCI also significantly reduced monocyte and neutrophil entry into the SCI lesion (Campbell *et al*., 2008). Thus, the liver appears to function as a “biological rheostat” that titers neuroinflammation, suggesting that it may be a clinically feasible target for modulating inflammatory-mediated central nervous system (CNS) injury or repair.

Consistent with acute hepatic inflammation provoking CNS inflammation, we hypothesized that concomitant liver inflammation at the time of SCI would worsen spinal cord pathology and outcomes. This idea has clinical relevance as there are several groups of people at risk of hepatic inflammation at the time of SCI. One population is those with obesity. A common cause of liver inflammation is obesity-associated non-alcoholic fatty liver disease (NAFLD), which includes non-alcoholic steatohepatitis (NASH) (Z. Younossi *et al*., 2018; Polyzos, Kountouras and Mantzoros, 2019). NALFD and NASH have been on the rise in the United States in the last 20 years (Ng *et al*., 2014; Rinella, 2015). Not surprisingly, ∼25% of people who sustain SCI are obese (Stenson et al., 2011) and are likely to have underlying hepatic inflammation at the time of injury. Similarly, those with a history of alcohol or drug abuse sustaining SCI have a high probability of hepatic inflammation at the time of injury (Garrison *et al*., 2004). A third “at-risk” group with potential pre-existing hepatic inflammation is the aged population, whose incidence of SCI has been steadily increasing in recent years (Rivara *et al*., 1993; Garrison *et al*., 2004; Chen, He and DeVivo, 2016; Hunt *et al*., 2019). Thus, a substantial number of spinal cord injuries may occur on a background of hepatic inflammation, making it critical to understand how an inflamed liver at the time of injury affects long-term intraspinal, metabolic and functional outcomes.

To test our hypothesis that pre-existing liver inflammation worsens outcomes from SCI, we used a transient “gain-of-function” surgical cholestasis model to induce liver inflammation in rats just prior to SCI. Our results show that pre-existing hepatic inflammation at the time of SCI exacerbates locomotor impairments, worsens spinal and hepatic pathology, and enhances metabolic disruption induced by SCI. We also detected a significant increase in Fetuin-A mRNA in the liver and Fetuin-A protein in the serum and the cerebrospinal fluid (CSF) after SCI, implicating it as a mediator of tissue pathology at the injury site. Indeed, when recombinant Fetuin-A was microinjected into the intact spinal cord, this liver-derived protein elicited neuroinflammation and significant neuron loss. Thus, pre-existing liver inflammation significantly worsens the intraspinal and systemic consequences of SCI, and Fetuin-A may be a novel liver-derived mediator of neuropathology. Overall, managing acute liver inflammation may be a novel therapeutic target for improving recovery and metabolic health after SCI.

## METHODS

All procedures were approved by the Institutional Animal Care and Use Committee (IACUC) and followed standards established by the NIH and The Ohio State University animal care guidelines. Female Sprague Dawley rats (∼250g; Harlan Laboratory) were used for all studies. All rats were randomly assigned to treatment groups and all outcome measures were assessed in a blinded manner. Rats were housed in a climate-controlled facility (22-24°C) in ventilated cages with a 12-hour light/dark cycle with lights on between 0700 and 1900 local time. *Ad libitum* access to water was provided in each cage. All rats were group-housed up to 3 rats per cage. Rats were fed a nutritionally complete diet (Teklad Global 16% Protein Rodent Diet) consisting of 66% energy from carbohydrate, 22% from protein and 12% from fat.

### Bile Duct Ligation (BDL)

Bile duct ligation (BDL) is commonly used as a model of biliary acid accumulation and hepatic inflammation (Kountouras, Billing and Scheuer, 1984; Marques *et al*., 2012). Here this model was used since it is simple and reliable and could easily be adapted to our SCI model. To determine when significant hepatic inflammation occurs after BDL, pilot studies were conducted. Rats were anesthetized by intraperitoneal ketamine (80 mg/kg) and xylazine (10 mg/kg). After laparotomy, the common bile duct was exposed at the level of the duodenum. Two methods were used to ligate the common bile duct. The first, which was used for all experiments referred to as “permanent BDL,” used two nylon 4-0 sutures spaced 0.5 mm apart around the common bile duct to occlude bile flow. The second method, which was used for all experiments referred to “reversible BDL” or “rBDL”, used a single strand of 0.5mm wide umbilical tape tied around the common bile duct. The thick umbilical tape obstructed bile flow while also allowing for subsequent easy removal and restoration of bile flow, which is not possible with sutures. Sham surgeries involved laparotomy and bile duct exposure but no ligation. The laparotomy was closed with a continuous suture and animals were injected with 10cc of saline and 0.25cc of 0.05 mg/kg buprenorphine subcutaneously for pain relief. Buprenorphine can have inflammatory or anti-inflammatory effects on different tissues and systems, including the liver (Hervé *et al*., 2004; Blaha and Leon, 2008; Guarnieri *et al*., 2012). Thus, all rats receiving a laparotomy were administered buprenorphine for 5 days to eliminate any confounds from drug-induced effects on hepatic inflammation. Animal weights were monitored daily and daily injections of saline were given to maintain hydration. At 1d or 5d post-ligation or post-sham surgery (n=5/group), rats were anesthetized with 1.5X surgery dose of ketamine/xylazine and exsanguinated via intracardiac perfusion with DEPC-treated phosphate buffered saline (PBS). A 0.5g segment of fresh liver was collected from the right medial lobe and flash frozen in liquid nitrogen. Total fresh liver weight and body weight were noted. This study revealed consistent and significant hepatic inflammation by 5d compared to time-matched sham controls, so in subsequent studies, SCI was induced at 5d post-BDL or 5d post-rBDL.

### Moderate Midthoracic Spinal Cord Contusion Injury

Briefly, 5d after BDL or rBDL, a laminectomy was performed at T8 and the exposed spinal cord was contused using the Infinite Horizons device (Precision Systems and Instrumentation) at 200 kD (Tripathi and McTigue, 2008; Sauerbeck *et al*., 2013). Back muscles were sutured, and the skin was closed with wound clips. Twice daily manual bladder expressions were performed until spontaneous voiding returned (∼14d post-injury).

For rats that received reversible BDL+SCI (rBDL+SCI), just prior to SCI the laparotomy was re-opened and and the umbilical tape around the bile duct was carefully removed with micro-scissors to restore bile flow. The abdominal cavity was sutured and rats proceeded to SCI surgery.

### BDL + SCI Experimental Groups

In the permanent BDL study, rats were randomized into the following groups: laparotomy sham (n=5), BDL (n=5), SCI (received sham BDL surgery; n=8) and BDL+SCI (n=5). Rats from all three experiments were sacrificed at 28 days post-ligation. Five rats in the permanent BDL+SCI died from either a ruptured bile duct (n=2) or ascites accumulation (n=3) ∼20d post-SCI. All irreversible BDL and BDL+SCI rats exhibited signs of jaundice and changes in urine acidity by the endpoint of the study.

### rBDL + SCI Experimental Groups

In the rBDL study, rats were randomized to the following groups: laparotomy sham (n=4), rBDL (n=4), SCI (received sham rBDL surgery; n=7), and rBDL+SCI (n=7). A subsequent independent replication rBDL experiment was performed with sham (n=3), rBDL (n=3), SCI (n=6), rBDL+SCI (n=6) groups. Rats were sacrificed at 28 days post-ligation/23 days post-ligation reversal + SCI. A separate cohort of rats was randomized into the following groups: sham (n=5), rBDL (n=5), SCI (received sham rBDL surgery; n=5), and rBDL+SCI (n=5). Rats from this cohort were sacrificed at 6 days post-ligation/1-day post-ligation reversal + SCI. No pre-mature mortality occurred with any of the rBDL+SCI study groups. No rats that received either rBDL alone or rBDL+SCI exhibited signs of jaundice or changes in urine acidity.

### Open Field Activity Assessment - Activity Box

Three-dimensional movements of individual animals in an open-field environment were measured using an activity monitoring device (Opto-M3, Columbus Instruments, Columbus, OH) as previously described (Popovich *et al*., 2014). Animals were placed in activity boxes for 30 minutes. Data for each 30-minute period was divided into three 10-minute segments and average. Animals were tested pre-BDL then at 1-, 2- and 3-weeks post-SCI.

### Hindlimb Locomotor Assessment

All rats were acclimated to handling and testing in the open field environment prior to any surgeries. Rats were tested by two observers blinded to the treatment groups at 1 and 5 days post-BDL and on days 1, 3, 7, 10, 14 and 21 post-SCI using the Basso-Beattie-Bresnahan (BBB) locomotor rating scale (Basso, Beattie and Bresnahan, 1995). Hindlimb scores for each rat were averaged and used to create group means at each day. BBB subscores were calculated and reported as previously described (McTigue *et al*., 2007). If BBB and activity box testing fell on the same day, activity box testing occurred at least 2h after BBB testing.

### Tail Vein Blood Collection

Lateral tail vein blood samples were collected every 4-5 days. Rats were anesthetized with 5% isoflurane and tails were immersed in warm water. A 28-gauge butterfly needle was inserted into one of the veins and ∼500 μl of whole blood was collected and centrifuged for 10 min at 300g under sterile conditions to separate serum and plasma. ∼200 μl of serum and plasma were collected from each rat at each time point. Flash-frozen aliquots of serum and plasma were stored at -80OC until analysis.

### CSF Collection

CSF from the cisterna magna was collected just after tail vein blood was withdrawn. Rats were affixed to a stereotaxic frame; a midline incision was made at the nape and the underlying muscles were cut and separated to expose the cisterna magna. A pulled glass capillary with an ∼40µm beveled tip was inserted into the dura and ∼50 μl of CSF was collected with a butterfly needle and 1 ml syringe under sterile conditions without blood contamination. Samples were flash frozen and stored at -80oC until analysis. The muscles were sutured, and the incision closed with surgical staples. Animals were returned to their home cages and allowed to recover on heated pads until fully ambulatory.

### Fetuin-A intraspinal microinjections

Recombinant Fetuin-A (Sigma; SRP6217) was diluted in 0.9% saline vehicle solution. All solutions were filter sterilized using a 0.22 μm filter and mixed immediately before use. Adult female Sprague-Dawley rats (∼ 250 g; n = 27) were randomly assigned to treatment groups (0.9% saline vehicle control, 10 μM Fetuin-A and 100 μM Fetuin-A (n = 8/group), then anesthetized for surgery as above. Using aseptic technique, a laminectomy was performed at the T8 vertebral level. Custom pulled UV-sterilized glass micropipettes beveled to an outer tip diameter of 25–40 μm were loaded with the proper solution. Two independent microinjection experiments using a hydraulic micropositioner were completed (David Kopf Instruments, Tujunga, CA). The first group received a unilateral microinjection into the white matter/gray matter border (n=3/group) using coordinates 0.7 mm lateral and 1.1mm ventral to the dorsal spinal cord midline. The second was unilaterally injected into the ventral gray matter (n=5/group) using coordinates 0.5 mm lateral and 1.2 mm ventral to the dorsal spinal cord midline. A 500 nl bolus injection was slowly administered using a PicoPump (World Precision Instruments). Injection sites were labeled with sterile charcoal (Sigma), muscles surrounding the laminectomy were sutured, skin was stapled with wound clips, and rats were given 5 ml sterile saline subcutaneously before being placed into a heated recovery cage. Rats were sacrificed and transcardially perfused with 4% PFA 3 days post-injection as described earlier. Spinal cords were sectioned at 10 µm and collected sections spanned the entire rostral caudal extent of the injected area as previously described (Goldstein *et al*., 2017).

### Tissue Extraction and Preparation

At the end of each SCI study, rats were fasted for 6 hours, then anesthetized with 1.5X surgical dose of ketamine/xylazine. Cardiac blood samples were collected and centrifuged for 10 min at 300g. Serum was removed, flash frozen and stored at -80oC until analysis. Next, rats were exsanguinated with DEPC PBS and a 0.5g segment of fresh liver was collected from the right medial lobe and flash frozen in liquid nitrogen. Next, rats were perfused with 4% paraformaldehyde (PFA) in 0.1 M PBS. Liver, brain, spinal cord, mesenteric fat and retroperitoneal fat were removed and post-fixed in 4% PFA for 2 h and transferred to 0.2 M phosphate buffer (PB) overnight. Total fixed liver, fat and body weights were recorded. Samples then were removed from PB and cryoprotected for 2-3 d in 30% sucrose solution. Fixed liver, brain and spinal cord samples were quickly frozen on dry ice and blocked using optimal cutting temperature (OCT) solution. 10 μm thick spinal cord and 20 μm thick brain and liver sections were sectioned on cryostat at -20°C and slide-mounted (Superfrost Plus Slides, Fisher Scientific). Slides were stored at -20°C until used.

### Serum, Plasma and CSF Alanine Aminotransferase (ALT), Bile Salts, Ammonia, Endotoxin, Glucose, Insulin and non-esterified fatty acid (NEFA) Analysis

Serum ALT (Point Scientific, A7526), bile salts (Cell Bio Labs, STA-631), ammonia (Sigma-Aldrich, AA0100) and endotoxin (Thermo Scientific Pierce, 88282) were measured using spectrophotometric clinical assays at 450 nm with a microplate reader (ELx 800 uv; BIO-TEK Instruments Inc, Winooski, Vt) in accordance with each manufacturer’s instructions. Serum glucose was measured using a clinical assay in accordance with the manufacturer’s instructions (Point Scientific) at a wavelength of 340 nm using a spectrophotometer (DU 600; Beckman Coulter, Inc., Brea, CA, USA). Serum insulin was measured using a commercial kit (Millipore Corporation, Billerica, Mass) by an enzyme immunoassay at 450 nm with the aforementioned microplate reader. Serum NEFA were measured spectrophotometrically using a commercially available assay (Wako Diagnostics), as previously described (Li *et al*., 2017).

### Serum and CSF Fetuin-A Analysis

Serum and CSF Fetuin-A was measured with an ELISA kit (ELISA Kit, BD ELISA MBS019821, BD Pharmingen, San Diego, CA) according to manufacturer’s directions.

### Liver Histology

Immunohistochemistry was performed on liver tissue as previously described (Sauerbeck *et al*., 2015; Goodus *et al*., 2018). Briefly, anti-CD11b (Ox42; Serotec, MCA275, 1:2,000) and anti-Clec4f (ThermoFisher Scientific, PA5-47396, 1:400) were used to visualize liver macrophages (Kupffer cells). Liver non-heme iron was visualized using Perls Prussian Blue reagent and H-Ferritin (Santa Cruz, sc-376594, 1:1,000) as previously described (Schonberg and McTigue, 2009; Schonberg *et al*., 2012). Oil Red O histology (Sigma, cat# O0625) was used to assess lipid accumulation as previously described (Sauerbeck *et al*., 2015). Hematoxylin and eosin was used to visualize hepatocyte ballooning, inflammatory infiltrates and lipid deposition in the liver as previously described (Bruno *et al*., 2008; Masterjohn and Bruno, 2012).

### Brain and Spinal Cord Histology

20 μm thick coronal cross-sections of the brain were used to visualize microgliosis and astrocytosis in the cortical gray matter as previously described (Goodus *et al*., 2016; Morrison *et al*., 2017). Briefly, Iba1 (Wako, 019-19741, 1:2,000) was used to visualize microglia/macrophages and anti-GFAP (Dako, Z0334, 1:15,000) was used to visualize astrocytes. Cross-sections of the spinal cord spanning the rostral to caudal extent of the lesion and microinjection sites were used for immunohistochemistry as previously described (Sauerbeck *et al*., 2013; Goodus *et al*., 2018). Briefly, axons and white matter were visualized with anti-neurofilament (NF) antibody (DSHB, RT97, 1: 2000) and eriochrome cyanine (EC), respectively. Anti-CD11b (1: 2000) and anti-CD68 antibodies (Serotec, MCA341GA, 1:1000) were used to visualize macrophages/microglia. Non-heme iron was visualized using Perls Prussian Blue reagent as previously described (Schonberg and McTigue, 2009; Schonberg *et al*., 2012). Neurons were visualized with anti-NeuN antibody (EMD-Millipore, MAB377, 1: 25,000).

### Tissue Analysis and Quantification

Immunolabeled tissue sections were imaged at 16X magnification using an Axioplan 2 imaging microscope equipped with an AxioCam digital camera and AxioVision v.4.8.2 software (Carl Zeiss Microscopy GmbH, Jena, Germany) An investigator blinded to experimental groups performed all histological analyses using either MCID elite imaging software (Imaging Research Inc, https://mcid.co.uk.), Image J (NIH, https://imagej.nih.gov/ij/), or MIPAR image analysis software (https://www.mipar.us/). Liver sections were randomly selected from each group. The proportional area of Clec4f, CD11b, Oil Red O and Perls Prussian Blue stain was quantified from 6 random fields per section by dividing the labeled “target” area by the total hepatocyte area measured in each field. Staining levels are expressed as % of total hepatocyte area. Livers stained with hematoxylin and eosin were scored for signs of NAFLD and NASH as previously described (Bruno *et al*., 2008; Masterjohn and Bruno, 2012).

The density of Iba1+ microglia/macrophages and GFAP+ astrocytes in brain sections was quantified and averaged from 5 coronal brain sections spaced 100 μm apart in each brain as previously described (Goodus *et al*., 2016; Morrison *et al*., 2017). The proportional area of Iba1 and GFAP was quantified in a 200 μm x 200 μm region in the cortical gray matter as depicted in Supplemental figures 1F, 1H, 2F and 2H using MIPAR imaging software.

**Figure 1.**
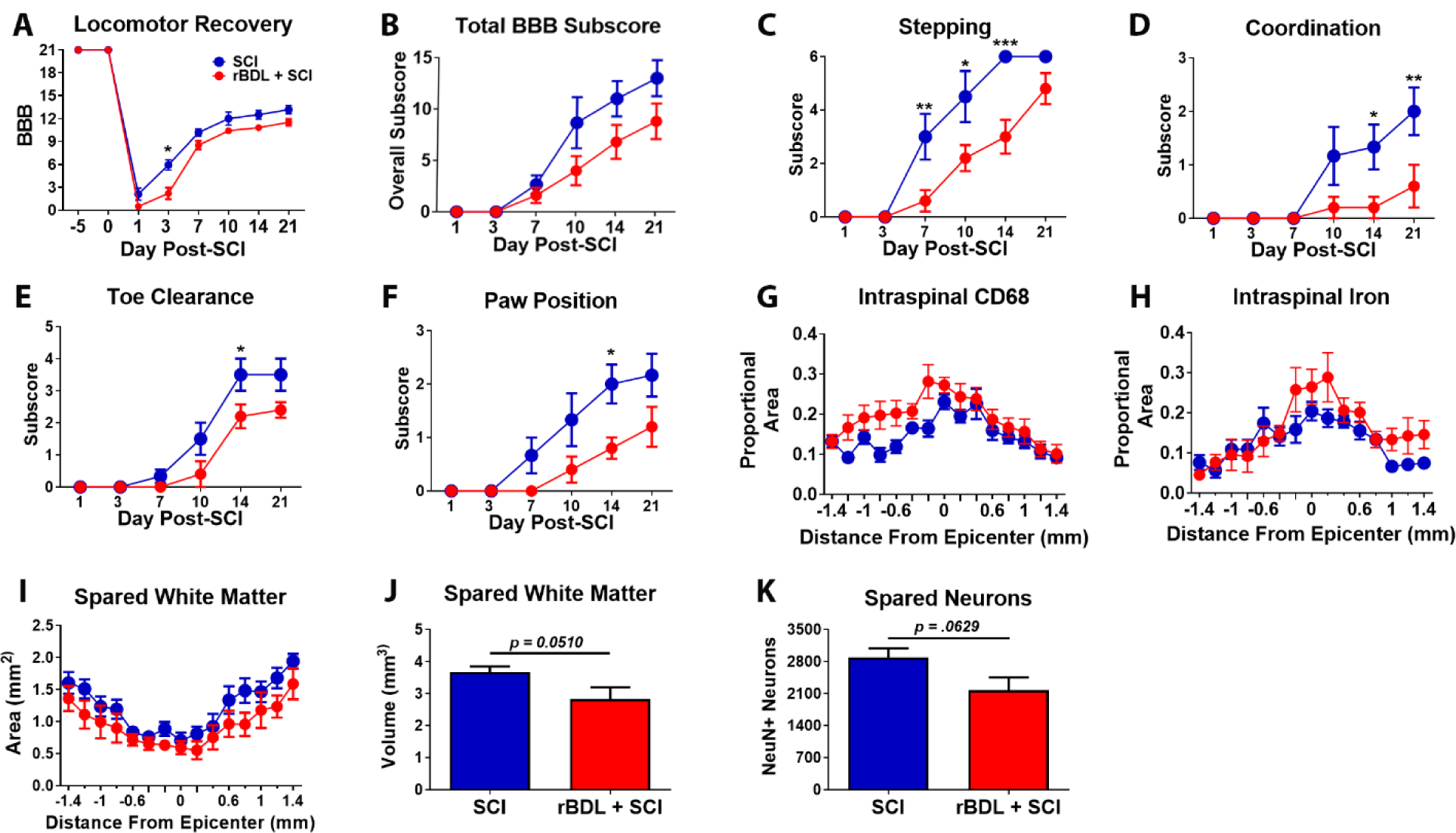
rBDL+SCI exacerbates locomotor deficits and increases intraspinal inflammation and iron accumulation. (A-F) Locomotor recovery and BBB subscores of rBDL+SCI rats were significantly lower than SCI rats. *p <0.05 and **p<0.01via RM ANOVA and Bonferroni post hoc test. Intraspinal (G) CD68 (p = .0486 RM ANOVA) immunolabeling and (H) iron (p = .0490 RM ANOVA) were increased in rBDL+SCI rats vs. SCI. (I) rBDL+SCI sections had reduced spared white matter area (p<0.001, treatment effect, RM ANOVA). (J, K) rBDL+SCI spinal cords showed trends for decreased white matter volume (p = .0510, student’s t-test) and spared neurons (p = .0629, student’s t-test). SCI, rBDL+SCI, n=7. Error bars represent ± SEM.

**Figure 2.**
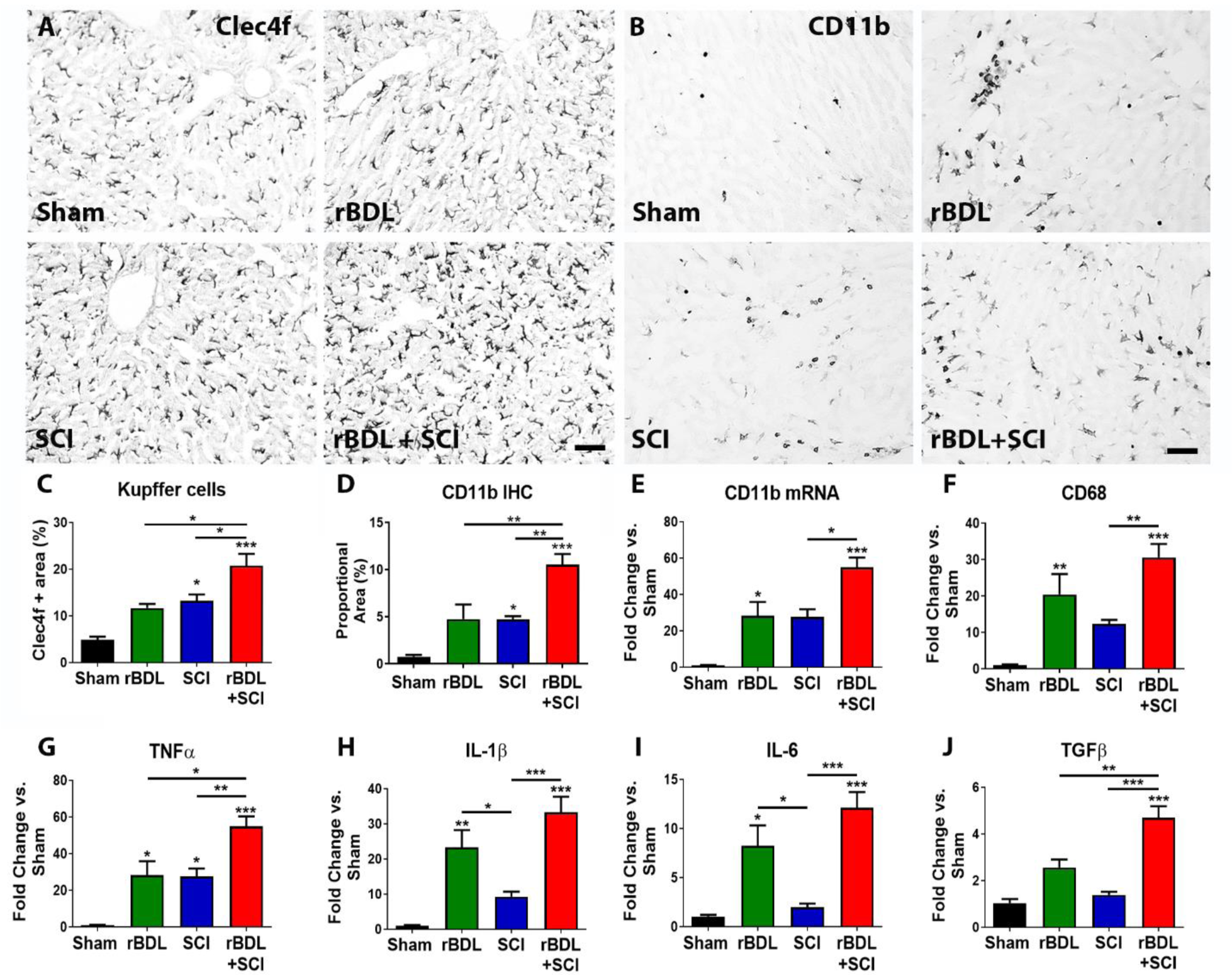
Increased hepatic inflammation at 23 dpi after rBDL+SCI. Representative images of (A) Clec4F and (B) CD11b stained livers at 28d post-ligation and 23d post rBDL+SCI show exacerbated Kupffer cell activation and inflammatory infiltrates in rBDL+SCI livers. Quantification of (C) Clec4f+ and (D) CD11b+ staining area expressed as % of total hepatocyte area showed significantly increased levels in the rBDL+SCI group compared to all other groups; both were also significantly higher in the SCI group vs. sham controls. Liver mRNA for (E) CD11b, (F) CD68, (G) TNFα, (H) IL-1β, (I) IL-6 and (J) TGFβ were elevated when SCI was performed on a background of liver inflammation. All genes were significantly increased in the rBDL+SCI rats compared to SCI. *p<0.05, **p<0.01 and ***p<0.001 via one-way ANOVA and Bonferroni post hoc test. Asterisks directly above bars represent significance vs. Sham. Sham and rBDL, n = 4; SCI and rBDL+SCI, n=7. Scale bars = 100 μm. Error bars represent +SEM.

For spinal cords, the injury epicenter was identified as the section with the least EC/NF labeling; analyses of spared white matter at the lesion epicenter and in 10 sections spaced 200 μm apart for a total of 2 mm analyzed rostral and caudal to the epicenter were completed using Cavalieri estimation (McTigue *et al*., 2007). The total immunoreactivities for CD68+ macrophages/microglia and Perls Prussian Blue iron label were quantified and converted to proportional area of the intact spinal cord tissue using MCID and ImageJ software at the lesion epicenter and in 10 sections spaced 200 μm apart for a total of 2 mm analyzed rostral and caudal to the epicenter. Areas of missing tissue were excluded from all analyses.

For Fetuin-A microinjected spinal cords, CD11b+ microglia/macrophages, NeuN+ neurons and Perls iron images were digitized and thresholded using MCID and/or ImageJ analysis software. Sample boxes (0.0961 mm2) were placed in the lateral white matter/gray matter border and ventral gray matter at both injection sites. The area of CD11b+ and Perls+ labeling were reported as a % of the total sample area in the white and gray matter. NeuN+ neurons were counted via MCID and/or ImageJ software.

### Analysis of Gene Expression

250 mg of frozen liver tissue was homogenized in Trizol Reagent (15596018, Life Technologies) to preserve nucleic acids. Samples were frozen at -80°C until used. RNA was isolated by a standard extraction protocol using chloroform/phenol (Almad and McTigue, 2010; Sauerbeck *et al*., 2015). The purity of all samples was assessed prior to amplification using microdrop spectrophotometry. The extracted RNA was synthesized into cDNA using Superscript III reverse transcriptase (Invitrogen) according to the manufacturer’s instructions. Expression of specific genes was assessed using quantitative Real Time Polymerase Chain Reaction (Applied Biosystems). Each qPCR reaction used 100 ng of cDNA with QuantiTec primers for tumor necrosis factor α (TNFα), CD68, CD11b, interleukin-1 alpha (IL-1α), interleukin-1 beta (IL-1β), CRP, Fetuin-A and SYBR green for detection (Table 1). The amplification of each sample was normalized using Quantum RNA18S (Applied Biosystems) gene expression as a control standard. The relative mRNA expression in each sample was determined with the ΔΔCT method and reported as fold-change vs. Sham controls (Schmittgen and Livak, 2008).

**Table 1.**
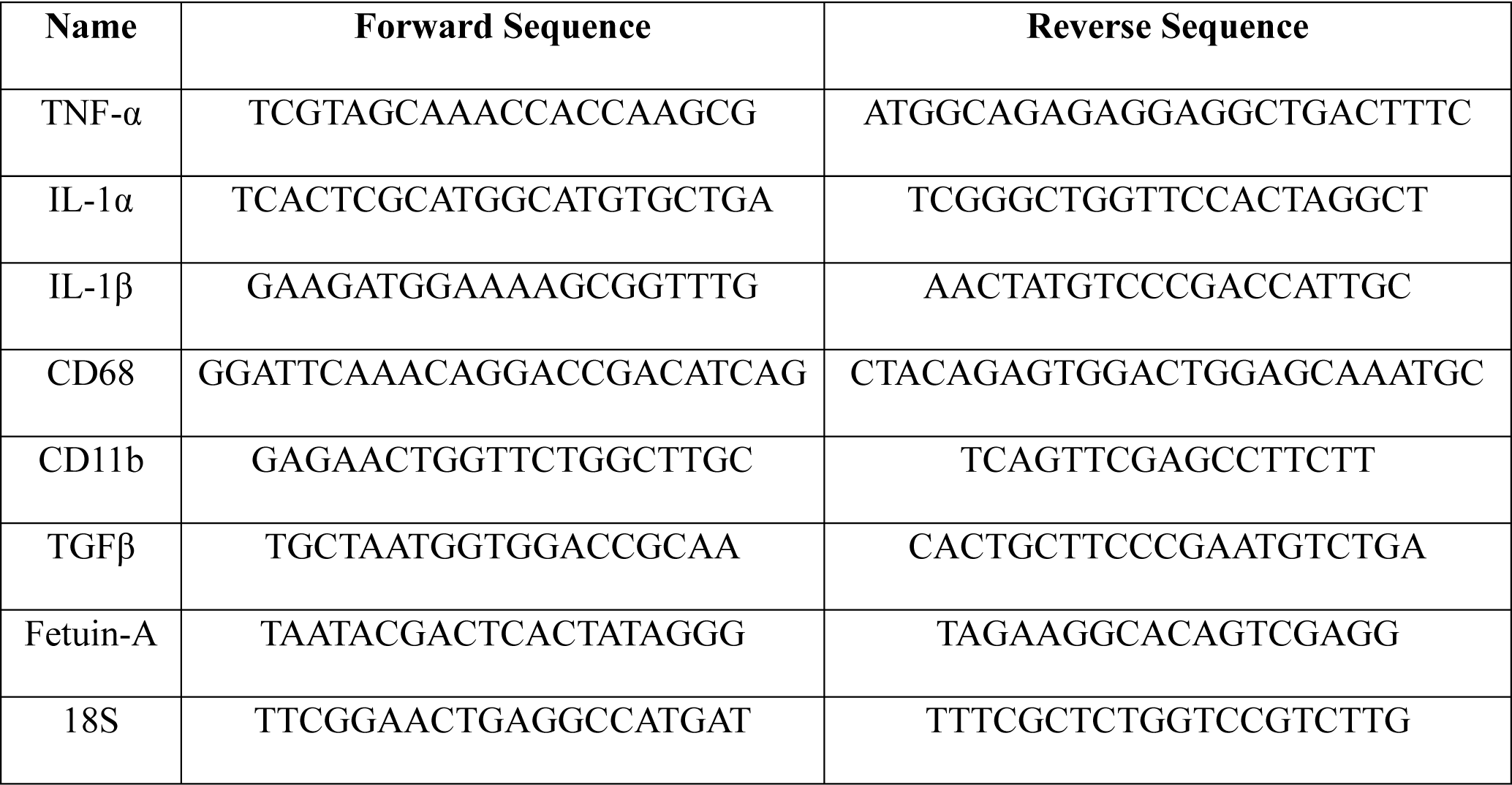
Primers for rtPCR quantification of hepatic gene expression in rats.

### Quantification and Statistical Analysis

Data (means ± SE) were analyzed using Graph Pad Prism 8.0 software. Student’s t-test, 1-way ANOVA or 2-way repeated measures ANOVA were used to examine main effects for all study endpoints when appropriate. Bonferroni post-hoc test was used to examine group differences following statistically significant main effects found via 1-way ANOVA; Bonferrroni post-hoc test was used to examine group differences via 2-way repeated measures ANOVA. Outliers that were greater than 2 times the standard deviation were identified for each data set via Grubb’s test and excluded from analysis. All analyses were considered statistically significant at p < 0.05. Homeostatic Model Assessment of Insulin Resistance (HOMA-IR) was calculated using the formula (Glucose concentration (mg/dl) x Insulin concentration (mU/L))/40.5 (Antunes *et al*., 2016).

## RESULTS

### SCI on a background of liver inflammation exacerbates intraspinal white matter loss, inflammation and iron accumulation and impairs locomotor recovery

To induce hepatic inflammation, we used a bile duct ligation (BDL) model to test the hypothesis that pre-existing inflammation exacerbates negative outcomes from SCI. While permanent BDL alone caused the expected liver inflammation 5d later (Fig. 1A, B), in rats surviving for 28d, it also caused sustained elevations in circulating bile salts and ammonia, which could confound results (Fig. 1C,D). Further, combining permanent BDL with SCI resulted in hepatic fibrosis and increased liver mass, and the animals displayed marked sickness behavior and brain gliosis reminiscent of hepatic encephalopathy at ∼3 weeks after SCI (Sup. Fig. 1C-J). These systemic comorbidities precluded a reliable analysis of the outcomes from SCI.

**Supplemental Figure 1.**
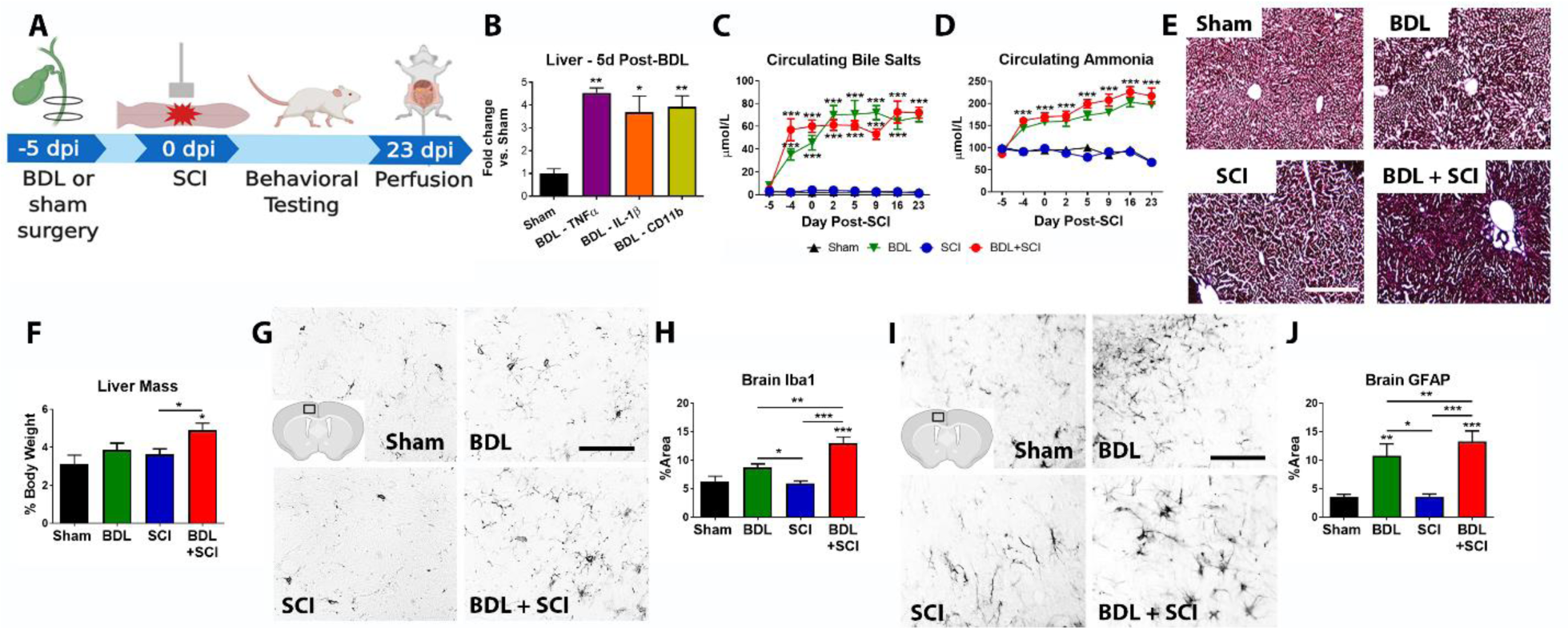
BDL+SCI Causes Liver Fibrosis and Brain Gliosis. (A) Experimental timeline. (B) Hepatic TNFα, IL-1β and CD11b were significantly elevated at 5d post-ligation compared to sham-operated controls. *p < 0.05 and ** p < 0.01 vs. Sham via ANOVA and Bonferroni post hoc test. n = 3/group. Circulating (C) bile salts and (D) ammonia were significantly elevated in rats that received BDL alone or BDL+SCI. **p < 0.01 and ***p < 0.001 vs. Sham and SCI via RM ANOVA and Bonferroni post hoc test. (E) Trichrome-stained livers from BDL+SCI rats showed extensive fibrosis and altered hepatocyte cytoarchitecture at 28d post-ligation and 23d post-SCI. (F) BDL+SCI liver mass was significantly increased vs. sham and SCI. *p <0.05 via ANOVA and Bonferroni post hoc test. Asterisk directly above bar represents significance vs. Sham. BDL+SCI rat brains showed significant (G, H) Iba1+ microglia activation and (I, J) GFAP+ astrogliosis at 23d post-injury. *p < 0.0, **p < 0.01and ***p < 0.001via ANOVA and Bonferroni post hoc test. Asterisks directly above bars represent significance vs. Sham. (C-J) Sham, BDL, BDL+SCI, n=5; SCI, n=8. Scale bars = 100 μm in D and 50 μm in F and H. Error bars represent ±SEM.

To overcome the complications of permanent BDL, we next tried releasing the ligated bile duct (rBDL) after 5d, a point at which consistent hepatic inflammation was observed (Sup. Fig. 2A,B). When these rats received SCI on the day of rBDL reversal, the confounds of chronically elevated bile salts and ammonia, hepatic fibrosis, sickness behavior and hepatic encephalopathy were avoided (Sup. Fig. 2C-J). Thus, our reversible rBDL model is a viable option for inducing SCI on a background of liver inflammation.

**Supplemental Figure 2.**
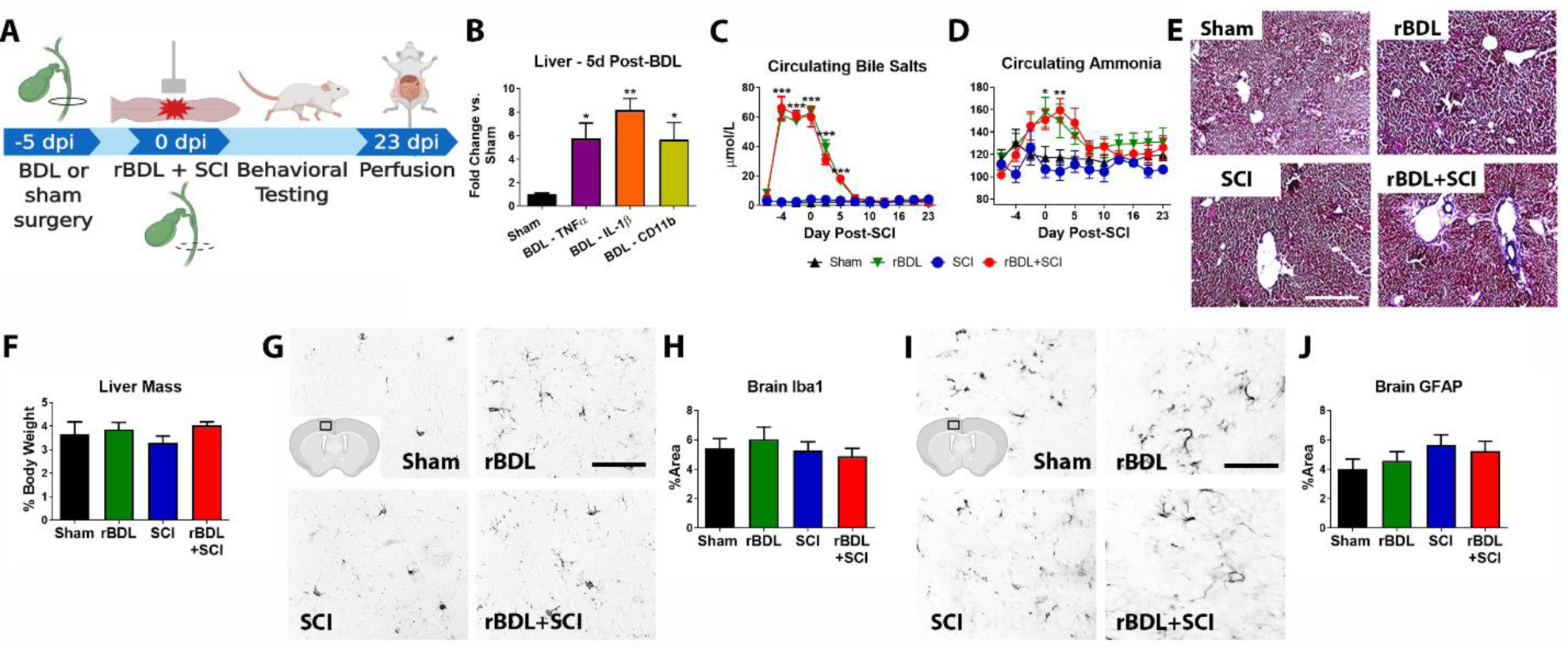
rBDL+SCI induces transient changes in circulating ammonia and bile salts, limited liver fibrosis and no signs of brain gliosis. (A) Experimental timeline. (B) Hepatic TNFα, IL-1β and CD11b mRNA were significantly elevated at 5d post-ligation compared to sham-operated controls. *p < 0.05 and ** p < 0.01 vs. Sham via ANOVA and Bonferroni post hoc test. n = 3/group. Circulating (C) bile salts and (D) ammonia were increased a day after BDL then returned to sham and SCI control levels 5d after ligation reversal. *p < 0.05, **p < 0.01 and ***p < 0.001 vs. Sham and SCI via RM ANOVA and Bonferroni post hoc test. (E) Trichrome-stained livers showed limited signs of fibrosis in rBDL+SCI rats at 28d post-ligation/23d post-SCI. (F) Liver mass was not different between groups (p = .3396, ANOVA). No significant differences in (G,H) Iba1+ microglia (p = .6729 ANOVA) or (I,J) GFAP+ astrocytes (p = .4266 ANOVA) were observed between groups. (C-J) Sham, rBDL, n = 4; rBDL, rBDL+SCI, n=7. Scale bars = 100 μm in D and 50 μm in F, H. Error bars represent ± SEM.

To test the hypothesis that liver inflammation exacerbates SCI-induced motor impairment, hindlimb locomotor recovery was examined in rats receiving rBDL+SCI vs. SCI only (plus sham BDL surgery). As expected, SCI only rats displayed an immediate loss of locomotor function that recovered to a plateau by ∼2w post-injury (Fig. 1A). Notably, rats receiving rBDL+SCI had significantly impaired hindlimb recovery compared to SCI alone (Fig. 1A,B). This was exhibited by reduced frequency of plantar stepping, reduced forelimb-hindlimb coordination, and fewer steps with toe clearance and parallel paw placement (Fig. 1C-F). Thus, motor recovery is significantly worse in rats with liver inflammation at the time of SCI compared to than those with normal livers.

We next examined if indices of lesion pathology in the spinal cord were altered by pre-existing hepatic inflammation, as would be suggested by the impaired functional recovery. First microglial and macrophage activity was examined in spinal cord cross-sections spanning the rostro-caudal extent of the injury using CD68 immunohistochemistry. Spinal cords from rBDL+SCI rats showed greater CD68+ macrophage/microglial accumulation compared with SCI alone rats, suggesting a greater level of microglial- or monocyte-derived macrophage activation and/or phagocytic activity in the rBDL+SCI spinal cords (Fig. 1G; *p=*.*0486 RM ANOVA, treatment effect*). Prior work showed that pathological levels of iron accumulate in SCI lesion sites following blood-brain barrier breakdown and cell death. Intraspinal iron is highly toxic and can cause macrophage activation. Thus, iron was quantified in the rostro-caudal extent of the lesions, which revealed that rBDL+SCI spinal cords had significantly more intraspinal iron compared with SCI alone (Fig. 1H; *p=*.*049 RM ANOVA, treatment effect*). Lastly, the amount of spared white matter and neuron numbers were quantified, which revealed significantly less spared white matter area in rBDL+SCI spinal cords across the extent of the lesion (Fig. 1I; *p<0*.*001RM ANOVA, treatment effect)*. This is consistent with other work showing that the amount of spared white matter correlates with the level of hindlimb recovery (Basso, Beattie and Bresnahan, 1995, 1996), both of which were worse here in rats receiving SCI on a background of hepatic inflammation. There also were trends for reduced overall white matter volume and neuron sparing compared to SCI controls (p=0.051 and p=0.0629, respectively; Fig. J, K). Collectively, these data show that rats receiving comparable SCIs have worse outcomes in terms of function and spinal lesions if the liver is inflamed at the time of SCI. Importantly, these data were independently replicated in a completely separate cohort of animals (Supp. Fig. 3).

**Figure 3.**
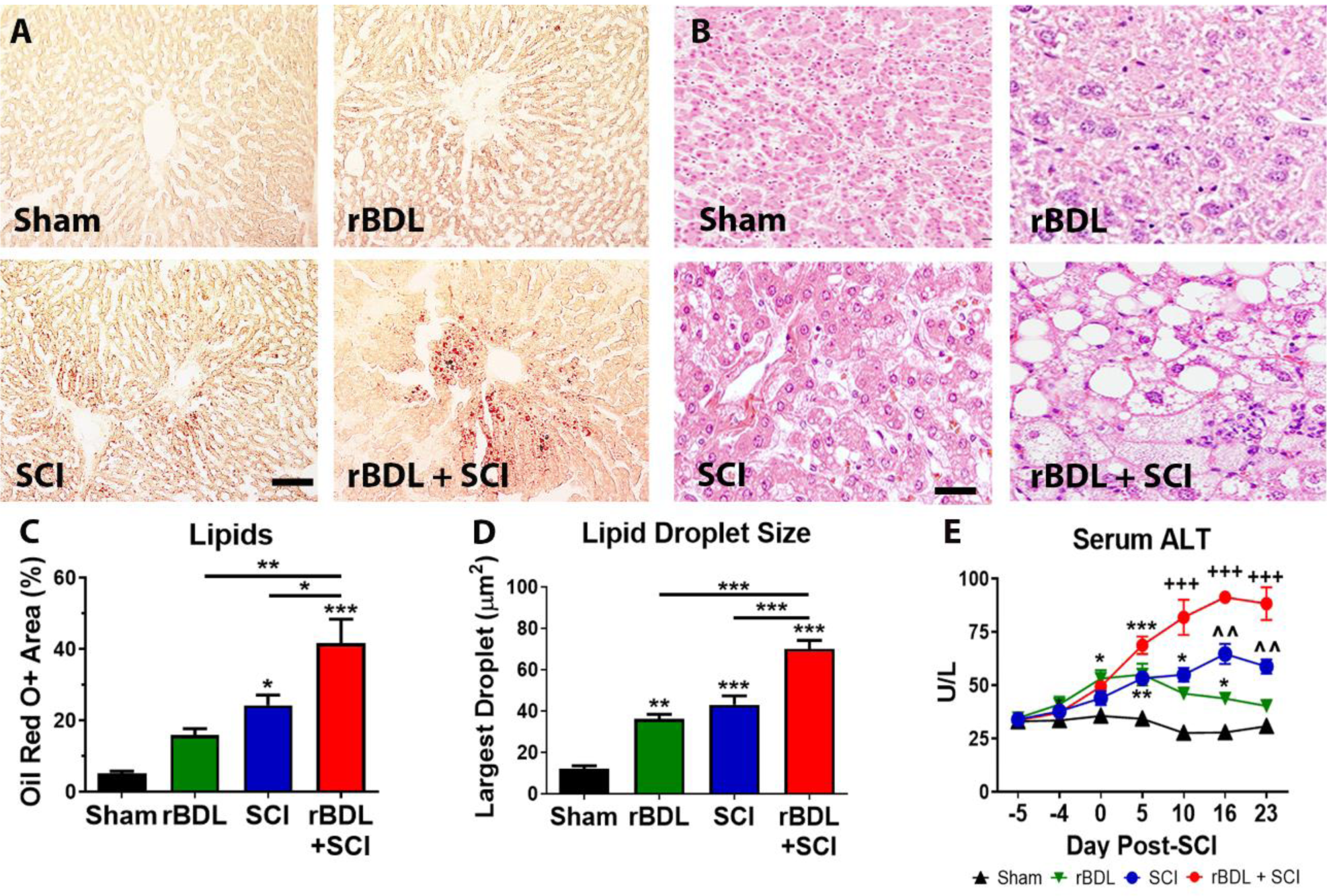
rBDL+SCI increases hepatic steatosis and hepatocyte damage. Representative images of (A) Oil red O and (B) hematoxylin and eosin stained livers at 28d post-ligation and 23d post-injury. SCI alone increases lipid accumulation, which is exacerbated in the rBDL+SCI livers, which show extensive hepatocyte ballooning. (C) Oil Red O+ staining area expressed as % of total hepatocyte area. (D) Quantification of largest fat droplet size revealed rBDL+SCI livers had the largest fat droplet size. *p<0.05, **p<0.01 and ***p<0.001 via one-way ANOVA and Bonferroni post hoc test in C and D. (E) Serum ALT levels transiently rose after rBDL and remained elevated in SCI and rBDL+SCI rats. rBDL+SCI doubled ALT compared to SCI alone. *p < 0.05, ** p< 0.01 and ***p < 0.001 vs. Sham, ^^ p < 0.01 vs rBDL and Sham and +++ p < 0.001 vs rBDL, SCI and Sham in (E) via RM ANOVA and Bonferroni post hoc test. Sham, rBDL n=4; SCI, rBDL+SCI, n = 7. Scale bars = 100 μm in (A) and 50 μm in (B). Error bars represent ±SEM.

**Supplemental Figure 3.**
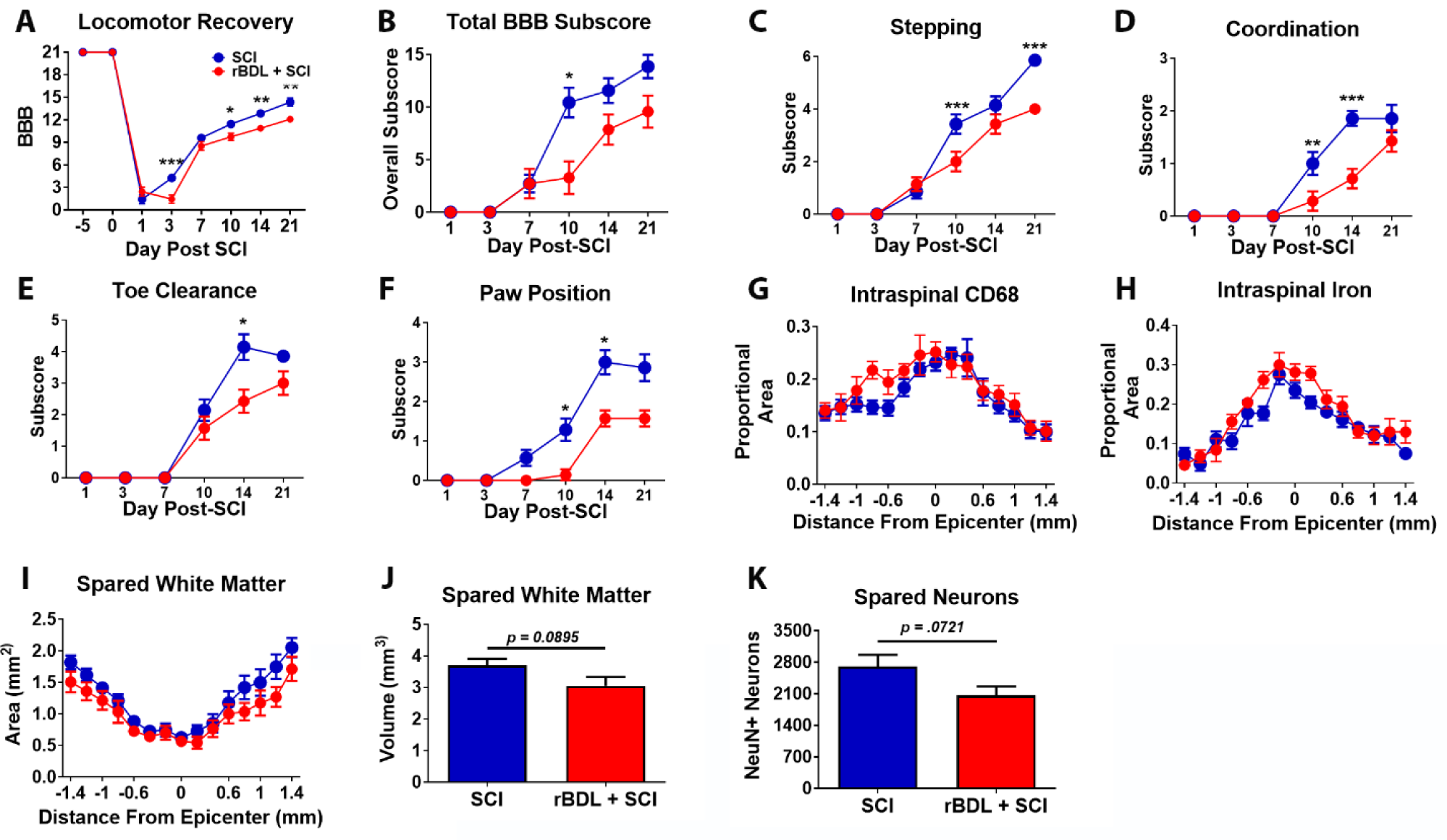
rBDL+SCI in a separate cohort of rats also exacerbates locomotor deficits and increases intraspinal inflammation and iron. (A-F) Locomotor recovery and BBB subscores of rBDL+SCI rats were significantly lower than SCI rats revealing greater motor impairments. *p <0.05, **p<0.01 and ***p<0.001in G via RM ANOVA and Bonferroni post hoc test. Intraspinal (G) CD68 (p = .0281 RM ANOVA) immunolabeling and (H) iron (p = .0009 RM ANOVA) were increased in rBDL+SCI rats vs. SCI. (I) rBDL+SCI sections had reduced spared white matter area (p<0.001, treatment effect, RM ANOVA). (J, K) rBDL+SCI spinal cords showed trends for decreased white matter volume (p = .0895, student’s t-test) and spared neurons (p = .0721, student’s t-test). SCI, rBDL+SCI, n=6. Error bars represent mean ± SEM.

### Post-SCI hepatic inflammation is exacerbated if the liver is inflamed at the time of injury

Our prior work showed that SCI alone causes NASH, including significant and prolonged liver inflammation and steatosis (Sauerbeck *et al*., 2015; Goodus *et al*., 2018). Given the elevated intraspinal pathology in rats receiving rBDL+SCI, we next determined if the combination of pre-existing liver inflammation plus SCI also exacerbated post-injury liver pathology. As expected, SCI alone significantly increased Kupffer cell number and activation, as seen by Clec4f and Cd11b immunoreactivity (Fig. 2A-D). Transient 5d rBDL also significantly increased Kupffer cell number and activation 23d later to a similar extent as SCI alone (although increases were not significant) (Fig. 2A-D). However, the combination of rBDL+SCI potentiated both measures compared to SCI or rBDL alone (Fig. 2A-D). Cd11b and CD68 mRNA were also measured as indices of Kupffer cell activation. Both were elevated 10-20-fold after SCI, although this was not significantly different from shams (Fig. 2E,F). Both also were significantly increased in the rBDL and rBDL+SCI groups compared to sham, and the rBDL+SCI levels were also greater than in SCI alone livers (Fig. 2E,F).

Increased hepatic cytokines typically accompany the progression of liver inflammation and are associated with stellate cell activation and fibrosis (Knittel *et al*., 1999; Gressner *et al*., 2002). Thus, several were measured here. As expected, SCI alone significantly increased TNFα mRNA expression ∼ 30-fold compared to shams; rBDL alone caused a similar rise in hepatic TNFα mRNA (Fig. 2G). Despite these large increases, the combination of rBDL+SCI potentiated hepatic TNFα mRNA >50-fold versus controls, which was significantly greater than in SCI or rBDL groups (Fig. 2G). IL-1β increased ∼9 -fold after SCI and significantly increased 20-30-fold in rBDL and rBDL+SCI livers, which were significantly greater than SCI or sham values (Fig. 2H). IL-6 mRNA only increased in rBDL and rBDL+SCI livers, suggesting its rise was mainly due to rBDL-induced signaling (Fig. 2I).

Lastly, TGFβ mRNA was examined as its expression is important for hepatic fibrosis. While neither SCI or rBDL altered TGFβ mRNA expression, the combination of rBDL+SCI significantly increased TGFβ mRNA compared to both groups and sham (Fig. 2J). Collectively, these data show that SCI-induced Kupffer cell activation, TNFα and TGFβ mRNA expression are all potentiated if the liver is inflamed at the time of injury. Further, even though fibrosis was not grossly visible in rBDL+SCI livers (Sup. Fig. 2E), the molecular signaling pathways triggered downstream of TNFα and TGFβ, which are involved in stellate cell activation and collagen deposition, appear to have been underway.

### SCI-induced hepatic fat accumulation and liver injury is exacerbated if the liver is inflamed at the time of injury

We previously reported that SCI in healthy rats induces rapid and prolonged hepatocyte lipid accumulation in rodents (Sauerbeck *et al*., 2015; Goodus *et al*., 2018). Here, we tested the hypothesis SCI-induced steatosis and hepatocyte injury will be exacerbated when SCI occurs on a background of liver inflammation. Using Oil red O (ORO) labeling and H&E staining, hepatocyte lipid accumulation was clearly visible in rats receiving SCI, and this lipid accumulation was significantly potentiated in rBDL+SCI rats (Fig. 3A,B). Quantification of confirmed a significant increase in ORO total area after SCI and in lipid droplet size in SCI and rBDL groups compared to sham (Fig. 3C,D). However, the combination of SCI plus rBDL synergistically increased both lipid amount and droplet size compared to SCI or rBDL alone (Fig. 3C,D).

Similarly, H&E labeling showed marked hepatocyte ballooning and cellular infiltrates in the rBDL+SCI livers compared to SCI or rBDL alone (Fig. 3B). These indices were quantified in H&E stained sections according to a previously established NASH scoring scale (Bruno *et al*., 2008; Masterjohn and Bruno, 2012), which confirmed increased histological hallmarks of steatosis and lobular and portal inflammation in rBDL+SCI livers compared to sham, rBDL alone or SCI alone (Table 2). Based on the evident hepatocyte pathology, serum ALT levels were measured, which we had previously shown increases after SCI (Sauerbeck *et al*., 2015; Goodus *et al*., 2018). Here again SCI alone induced a significant elevation in serum ALT by 10d post-injury, which was sustained through the end of the study (23d post-injury) (Fig. 3E). rBDL alone caused a transient rise in ALT that returned to baseline by 23d post-injury, mirroring the transient changes in serum bile salts and ammonia (Fig. 3E). The combination of rBDL+SCI significantly increased serum ALT levels compared to SCI or rBDL alone by 10d post-injury (Fig. 3E), which is consistent with the greatest hepatocyte ballooning seen in this group. These data further support the hypothesis that while liver pathology develops after SCI or rBDL, the pathology and damage to the liver (and spinal cord) is significantly greater if the liver is inflamed at the time of injury.

**Table 2.** Scoring criteria for identifying NASH in H&E stained liver sections. rBDL+SCI rats had increased signs of inflammation and lipid accumulation compared to all other experimental groups.

| Item | Definition | Score | rBDL Sham<br>n=4) | rBDL<br>(n=4) | SCI<br>(n=7) | rBDL + SCI<br>(n=7) |
| --- | --- | --- | --- | --- | --- | --- |
| **Steatosis** | (<5%) | 0 | 3 (100%) | 2 (66.7%) | 1 (14.3%) | 0.0% |
|  | (5-33%) | 1 | 0.0% | 1 (33.3%) | 5 (71.4%) | 0.0% |
|  | (>33-66%) | 2 | 0.0% | 0.0% | 1 (14.3%) | 3 (42.9%) |
|  | ( >66%) | 3 | 0.0% | 0.0% | 0.0% | 4 (57.1%) |
| **Fibrosis** | None | 0 | 3 (100%) | 3 (100%) | 7 (100%) | 6 (85.7%) |
|  | Perisinusoidal or<br>Periportal | 1 | 0.0% | 0.0% | 0.0% | 1 (14.3%) |
|  | Portal/periportal | 1 | 0.0% | 0.0% | 0.0% | 0.0% |
|  | Perisinusoidal<br>and<br>portal/periportal | 2 | 0.0% | 0.0% | 0.0% | 0.0% |
|  | Bridging fibrosis | 3 | 0.0% | 0.0% | 0.0% | 0.0% |
|  | Cirrhosis | 4 | 0.0% | 0.0% | 0.0% | 0.0% |
| **Ballooning** | None | 0 | 3 (100%) | 3 (100%) | 6 (85.7%) | 0.0% |
|  | Few | 1 | 0.0% | 0.0% | 1 (14.3%) | 2 (28.6%) |
|  | Many/Prominent | 2 | 0.0% | 0.0% | 0.0% | 5 (71.4%) |
| **Lobular**<br>**Inflammation** | No foci | 0 | 3 (100%) | 0.0% | 0.0% | 0.0% |
|  | <2 foci per 200X<br>field | 1 | 0.0% | 3 (100%) | 4 (57.1%) | 0.0% |
|  | 2-4 foci per<br>200X field | 2 | 0.0% | 0.0% | 3 (42.9%) | 5 (71.4%) |
|  | >4 foci per 200X<br>field | 3 | 0.0% | 0.0% | 0.0% | 2 (28.6%) |
| **Portal**<br>**Inflammation** | None to<br>Minimal | 0 | 3 (100%) | 1 (33.33%) | 4 (57.1%) | 0.0% |
|  | Greater than<br>Minimal | 1 | 0.0% | 2 (66.67%) | 3 (42.9%) | 7 (100%) |

### SCI concomitant with hepatic inflammation exacerbates SCI-induced increases in adipose tissue and insulin resistance

Data above show that SCI causes rapid development of NASH, which is exacerbated if the SCI occurs at a time that the liver is inflamed. NASH is the hepatic manifestation of metabolic disease and is often accompanied by increased abdominal obesity, hyperlipidemia, hyperglycemia and insulin resistance (Farrell and Larter, 2006; Park *et al*., 2012; Z. M. Younossi *et al*., 2018). Notably, these indices of metabolic dysfunction are often elevated in the SCI population as well (Kocina, 1997; Bauman and Spungen, 2001, 2008; Manns, McCubbin and Williams, 2005; LaVela *et al*., 2006; Weaver *et al*., 2007; Inskip *et al*., 2010; Rankin *et al*., 2017). In-house time course studies showed that NEFAs, glucose and insulin rise in a delayed fashion after SCI in rats, with significant peaks not present until 6 – 8 weeks post-injury (personal observation). The chronic rise was verified in a previous report that showed significant increases at 6w post-injury (Goodus *et al*., 2018). As expected, here the ∼3w post-SCI data show only slight non-significant changes in NEFAs, adipose, glucose and insulin, and no change at all in the rBDL group (Fig. 4). However, rBDL+SCI doubled the NEFA serum levels (significant main effect *p =* .*0457 one-way ANOVA*), and significantly increased mesenteric and visceral (not shown) adipose, serum glucose and insulin (Fig. 4A-D). Homeostatic Model Assessment of Insulin Resistance (HOMA-IR), which models insulin resistance using circulating glucose and insulin concentrations in humans and rodents (Antunes *et al*., 2016), was also significantly elevated in rBDL+SCI rats (Fig. 4E). Collectively, these data show that if the liver is inflamed at the time of SCI, SCI-induced liver pathology and metabolic dysfunction are accelerated and exacerbated.

**Figure 4.**
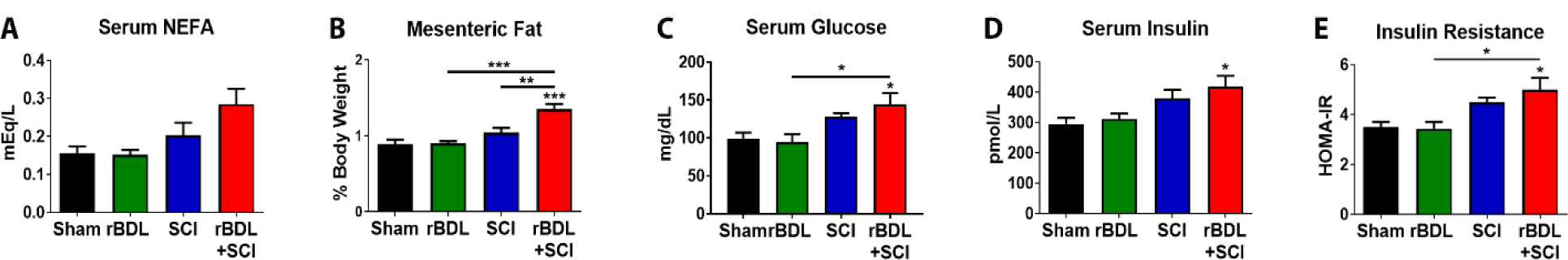
rBDL+SCI increases circulating fatty acids and white adipose tissue and disrupts circulating glucose and insulin homeostasis. rBDL+SCI increased (A) circulating NEFAs (p = .0457 ANOVA). (B) Mesenteric abdominal fat was significantly increased in rBDL+SCI rats compared to all other groups. rBDL+SCI increased serum (C) glucose, (D) insulin levels and (E) HOMA-IR values at 23dpi. *p<0.05, **p<0.01 and ***p<0.001 via one-way ANOVA and Bonferroni post hoc test. Asterisks directly above bars represent significance vs. Sham. Sham, rBDL, n=4; SCI, rBDL+SCI, n=7. Error bars represent +SEM.

### Hepatic iron and ferritin levels are increased in rBDL+SCI groups

Midthoracic SCI disrupts hepatic iron regulation and reduces circulating iron, which may contribute to the anemia and insulin resistance that occurs in rodents and humans after SCI (Anderson and Shah, 2013; Simcox and McClain, 2013; Britton, Subramaniam and Crawford, 2016; Goodus *et al*., 2018). Thus, hepatic iron levels and ferritin expression were compared in all groups. Perls labeling of non-heme iron was ∼2-fold higher in livers from SCI rats compared to sham (Fig. 5A,C). Iron accumulation was further exacerbated in livers of rBDL+SCI rats (Fig. 5A, C). Since iron accumulation stimulates ferritin expression, H-ferritin levels were also measured. Similar to iron, SCI alone increased H-ferritin ∼2-fold compared to sham controls, while H-ferritin was significantly elevated in the rBDL+SCI group (Fig. 5B, D). There was no significant difference between rBDL+SCI and SCI groups for iron or H-ferritin (Fig. 5C,D).

**Figure 5.**
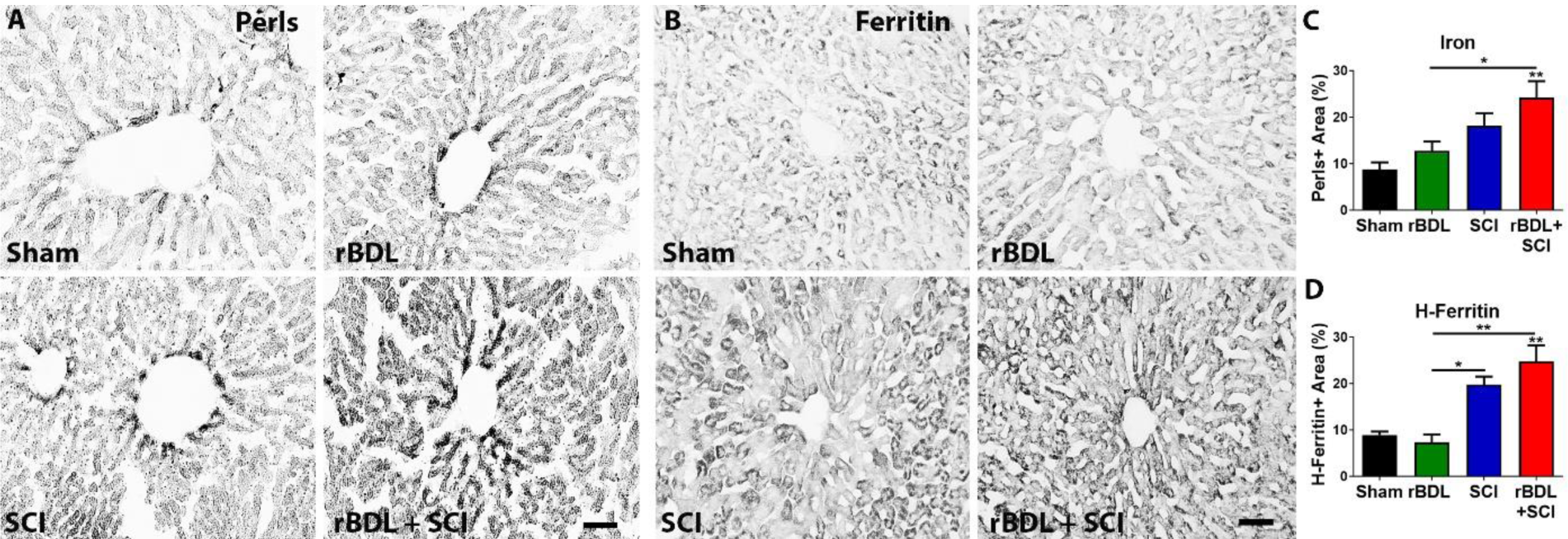
rBDL+SCI disrupts hepatic iron storage after SCI. Representative images of (A) Perls Prussian Blue stained liver sections and (B) liver H-ferritin immunolabeling at 28d post-ligation and 23d post rBDL+SCI show pre-existing liver inflammation increased liver iron stores after SCI. Quantification of (C) Perls+ and (D) H-ferritin+ staining area expressed as % of total hepatocyte area. Hepatic liver iron was elevated after rBDL+SCI compared to sham and rBDL alone, and liver ferritin was elevated in both rBDL+SCI and SCI groups compared to sham and rBDL controls. *p<0.05 and **p<0.01 via one-way ANOVA and Bonferroni post hoc test. Asterisks directly above bars represent significance vs. Sham. Sham, rBDL, n=4; SCI, rBDL+SCI, n = 7. Scale bars = 100 μm. Error bars represent +SEM.

### Liver inflammation at the time of SCI potentiates SCI-induced endotoxemia and increases hepatic, circulating and CSF levels of Fetuin-A

Our prior work and that of others shows that SCI in otherwise intact animals causes rapid gut dysbiosis and bacterial translocation, with bacteria detectable in the liver within 7d (Gungor *et al*., 2016; Kigerl *et al*., 2016; Kigerl, Mostacada and Popovich, 2018; O’Connor *et al*., 2018). Translocation of gut-derived bacteria and endotoxins (e.g., lipopolysaccharides) to the liver via the portal vein may cause or contribute to liver inflammation after SCI by activating immune receptors on hepatocytes, Kupffer cells and stellate cells (Goodus and McTigue, 2020). Here we measured serum endotoxin and again detected a significant increase after SCI, which occurred by 5d post-injury (data not shown) and remained significantly elevated through 23d post-injury (Fig. 6A). Bile duct ligation also has been associated with bacterial translocation (Deitch *et al*., 1990; Fouts *et al*., 2012). Here serum endotoxin was elevated ∼2-fold in this group throughout the study (not significant) (Fig. 6A). Similar to other measures above, rBDL+SCI had a synergistic effect on serum endotoxin levels, causing a significant increase by 23d post-injury compared to all other groups (Fig. 6A).

**Figure 6.**
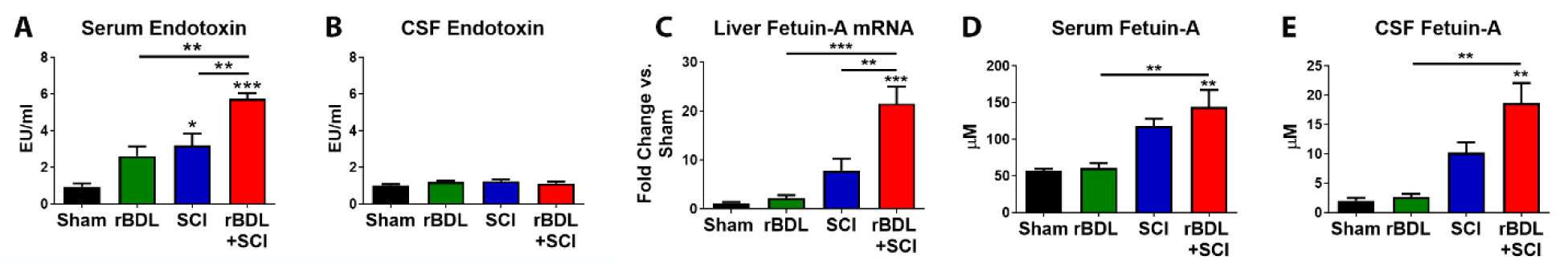
rBDL+SCI increases circulating endotoxins and Fetuin-A expression. (A) Serum endotoxin levels were significantly elevated in SCI and rBDL+SCI rats vs. sham and rBDL groups. (B) CSF endotoxin levels were unchanged in all groups. (C) Hepatic Fetuin-A mRNA was significantly increased in rBDL+SCI rats vs. shams at 23 dpi. (D) Circulating and (E) CSF Fetuin-A protein was elevated in rBDL+SCI rats compared to all other groups. *p < 0.05, **p < 0.01 and ***p <0.001 via one-way ANOVA and Bonferroni post hoc test. Asterisks directly above bars represent significance vs. Sham. Sham, rBDL, n = 4; SCI, rBDL+SCI, n = 7. Error bars represent +SEM.

Endotoxins can activate toll-like receptor 4 (TLR4) and this leads to significant neuropathology when it occurs within the CNS (Schonberg, Popovich and McTigue, 2007; Schonberg and McTigue, 2009; Kigerl *et al*., 2014; Goldstein *et al*., 2017). Because the blood-brain barrier is disrupted for several weeks after SCI (Popovich *et al*., 1996), it is possible that elevated circulating endotoxins could have entered the CSF in appreciable levels and enhanced spinal pathology in the rBDL+SCI. However, endotoxins were not different in the CSF between any groups (Fig. 6B), suggesting that endotoxins did not exacerbate intraspinal pathology in the rBDL+SCI group.

The liver can produce TLR4 ligands, such as the acute phase protein Fetuin-A, which is strongly associated with hepatic inflammation and lipid accumulation, obesity and diabetes (Pal *et al*., 2012; Trepanowski, Mey and Varady, 2015). Fetuin-A expression has not been examined after SCI, but if it is increased in the circulation and/or enters the CSF, it could be a potential mediator of exacerbated post-SCI spinal cord inflammation. Notably, SCI alone increased hepatic Fetuin-A mRNA ∼8-fold over sham control rats at 23dpi (Fig. 6C). While rBDL did not change Fetuin-A mRNA levels, the combination of rBDL+SCI significantly increased hepatic Fetuin-A mRNA ∼20-fold vs. shams, which was also significantly greater than in SCI and rBDL groups (Fig. 6C). Next, we examined Fetuin-A protein levels in the serum, which rose ∼2-fold in the SCI group and significantly increased 2.5-fold in rBDL+SCI rats (Fig. 6D). To determine if Fetuin-A may have accessed the injured spinal cord, CSF levels of Fetuin-A protein were measured, which revealed a ∼4-fold over shams in the SCI group and a significant ∼10-fold increase over sham in the rBDL+SCI group (Fig. 6E). Thus, liver inflammation at the time of SCI promotes Fetuin-A production, which increases Fetuin-A protein levels in the serum and CSF, revealing that the injured spinal is exposed to Fetuin-A after SCI+rBDL.

### Intraspinal Fetuin-A causes inflammation and neuron loss

To determine whether concentrations of Fetuin-A present in the CSF after SCI or rBDL+SCI, could exert biological effects on the spinal cord, recombinant Fetuin-A was microinjected into uninjured thoracic spinal cord white matter (WM) or gray matter (GM) at 10 μM, a concentration that approximates post-SCI levels in the CSF (see Fig. 6E). Additional groups examined the effects of a 10-fold a higher concentration (100 μM) of Fetuin-A and vehicle. At 3d post-injection, 10 μM Fetuin-A increased Cd11b+ microglia and macrophages 3 – 4-fold in WM and GM (not statistically significant), while 100 μM Fetuin-A significantly increased CD11b+ cells in both regions (Fig. 7A, D-F). Both Fetuin-A concentrations were neurotoxic; NeuN+ neurons were reduced ∼50-75% in GM injection sites compared to controls (Fig. 7B, G-I), revealing that the concentration of Fetuin-A in the CSF after SCI was at neurotoxic levels.

**Figure 7.**
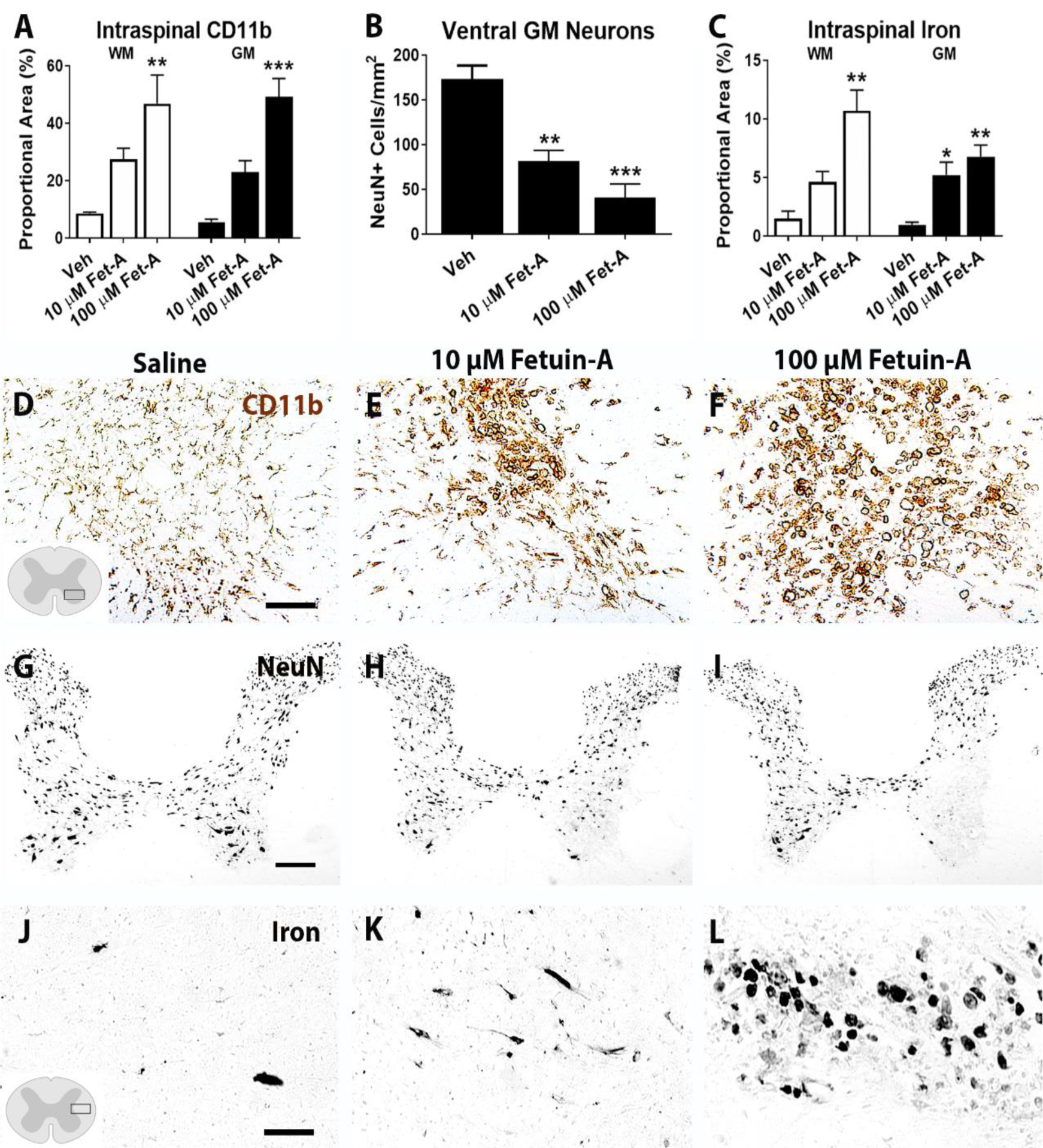
Intraspinal injection of Fetuin-A increases macrophages/microglia, reduces neurons and increases iron. Injection of recombinant Fetuin-A into the lateral white matter and ventral gray matter significantly increased (A, B, F, G, H) CD11b+ macrophages/microglia, reduced (C, I, J, K) NeuN+ neurons and increased (D, E, L, M, N) iron. Representative images in F, G, and H are from boxed region in diagram in F. Representative images in L, M and N are from boxed region in diagram in L. *p < 0.05, **p<0.01 and ***p<0.001vs. vehicle by one-way ANOVA and Bonferroni post hoc test. WM injections, n=3/group; GM injections, n=5 per group. Data represent mean ± SEM. Scale bars in (F) = 50 μm, (I) = 100 μm and (L) = 50 μm.

Work by our lab and others showed that intraspinal TLR4 activation causes iron accumulation, especially within macrophages (Zhang *et al*., 2005; Schonberg and McTigue, 2009; Goldstein *et al*., 2017) and that iron can mediate neuropathology (Goldstein *et al*., 2017). Thus, iron levels were also measured, which revealed that both Fetuin-A concentrations significantly increased iron levels in the GM, and the higher dose increased iron in the WM as well (Fig. 7C, J-L). Collectively, these novel data suggest that if the liver is inflamed at the time of SCI, liver-spinal cord axis disruption is exacerbated; this may be mediated in part through the promotion of Fetuin-A release that in turn could enhance chronic intraspinal pathology; it may also act locally in the liver to exacerbate post-SCI inflammation and iron accumulation.

## DISCUSSION

Results of this study show liver inflammation accentuates pathology and functional loss after SCI. While it is known that acute liver inflammation can enhance inflammatory cell infiltration into the inflamed CNS (Campbell 2005, 2008a, 2008b), the long-term effects of hepatic inflammation on CNS, hepatic or systemic pathology after CNS insult, however, were unknown. Our results report the striking finding that if an injury to the spinal cord occurs concurrent with liver inflammation, multiple long-term outcomes are worse, including intraspinal inflammation and tissue loss. This is critically important as the amount of tissue spared at the SCI site has a direct impact on extent of functional recovery attained after the injury (Basso et al., 2005, 2006). Consistent with that, rats with liver inflammation at the time of SCI had significantly worse walking abilities compared to control SCI rats. Further, rats with hepatic inflammation plus SCI had significantly more liver pathology than SCI controls, including greater Kupffer cell activation, more TNFα production, greater lipid accumulation and hepatocyte ballooning, more iron accumulation and higher levels of serum ALT. Indices of metabolic syndrome also occurred earlier in these rats. Our prior work showed that SCI causes the development of “neurogenic NASH” (Sauerbeck *et al*., 2015; Goodus *et al*., 2018), revealing that disrupting the spinal cord-liver axis causes extensive liver pathology. Here we show that perturbing the spinal cord-liver axis also damages the spinal cord and, depending on the severity of liver inflammation at the time of SCI, may even be detrimental to the brain. Notably, a clinical study of SCI subjects showed that blood analytes relating to “liver function” and “acute inflammation and liver function” including ALT were excellent predictors of neurological function at 3 and 12 months post-SCI (Brown *et al*., 2020). This suggests that liver pathology after SCI is an important biomarker and imply a two-way link between hepatic inflammation/injury and the outcome from SCI. Even in normal healthy animals, SCI evokes rapid liver inflammation, including Kupffer cell activation and hepatic cytokine production. Thus, based on the current results, combating the spontaneous liver inflammation that develops after SCI may provide a feasible approach for reducing pathology and dysfunction after injury to the spinal cord.

### Fetuin-A and TNFα: Potential mediators of a toxic spinal cord-liver axis after SCI that enhance intraspinal tissue damage and inflammation

Current results reveal the inflamed liver increases damage to the spinal cord after SCI, possibly through factors released into the circulation. While the specific factors are not yet known, several candidates exist. Here we identify Fetuin-A as potential hepatic-derived novel pathological mediator after SCI. Fetuin-A mRNA rapidly increased in the liver by 1d post-injury (not shown) and was sustained for 3 weeks. Fetuin-A protein also rose significantly in the serum and CSF of rBDL+SCI rats, indicating that the injured spinal tissue was exposed to elevated Fetuin-A in this group. Fetuin-A is a Toll-like receptor 4 (TLR4) agonist and prior work showed that intraspinal TLR4 activation by LPS or HMGB1 causes marked inflammation and cell loss (Schonberg, Popovich and McTigue, 2007; Schonberg and McTigue, 2009; Kigerl *et al*., 2018). Here we show Fetuin-A is similarly neurotoxic. After microinjection into intact spinal cords, Fetuin-A caused significant neuron death and robust microglia and macrophage activation. Thus, elevated Fetuin-A in CSF of rBDL+SCI rats could have mediated the increased intraspinal pathology in this group. A possible mechanism is through iron-mediated pathology. Elevated intraspinal iron is neurotoxic and intraspinal TLR4 activation increases local iron levels and causes neuron death (Schonberg and McTigue, 2009; Goldstein *et al*., 2017). Here intraspinal iron was elevated in both the rBDL+SCI rats and the Fetuin-A microinjection sites compared to controls. Thus, liver inflammation at the time of SCI enhances intraspinal macrophage and microglial activation and increases intraspinal iron, both of which could be potentially mediated, at least in part, by Fetuin-A.

Another potential liver-derived mediator of neurotoxicity is TNFα, as the current data show that pre-existing liver inflammation potentiates hepatic TNFα expression. If TNFα reaches spinal tissue at appreciable levels, it could directly stimulate intraspinal inflammation and potentially enhance other pathological processes, such as iron-mediated damage. For instance, TNFα can promote lactoferrin transport across the blood-brain barrier, which would increase intraspinal iron levels (Fillebeen *et al*., 1999). TNFα can also increase glutamate receptor density on neurons, thereby increasing their vulnerability to excitotoxicity (Ferguson *et al*., 2008), and it can promote an inflammatory macrophage phenotype within the spinal cord (Kroner *et al*., 2014). Considering that the blood-brain barrier remains leaky for at least one month after SCI (Popovich *et al*., 1996), exacerbated hepatic TNFα could be a potential mediator intraspinal inflammation and pathology.

As described above, liver inflammation may also increase the number of inflammatory cells that enter CNS sites of injury or inflammation. This could potentially be mediated by systemic TNFα. For instance, a study of brain inflammation found that the number of neutrophils entering the inflamed CNS were lower when animals received a systemic TNFα antagonist, showing that systemic TNFα can enhance inflammatory cell infiltration into the CNS (Campbell, 2007). Notably, a study of chronic SCI subjects revealed that >50% displayed elevated serum TNFα levels, revealing long-term peripheral inflammation, which could be derived, at least in part, from the liver (Hayes *et al*., 2002). A related way in which spinal cord pathology was increased could be due to increased mobilization of TNFα-producing monocytes to the spinal cord. Permanent BDL studies showed that at 10d after BDL, monocytes expressing TNFα infiltrated into the brains of the mice (Kerfoot et al., 2006). It is unclear, however, if this would have occurred in our model since we used transient BDL that only lasted 5 days.

### Liver damage after SCI is exacerbated if the liver is inflamed at the time of SCI

As stated above, liver inflammation may be an important biomarker for SCI severity (Brown *et al*., 2020). Our prior preclinical work shows that the liver is not only a “reporter” of injury but that the liver itself is also a “victim” of SCI, as it develops robust and persistent inflammation and lipid accumulation consistent with development of “neurogenic” NASH (Sauerbeck *et al*., 2015; Goodus *et al*., 2018; Goodus and McTigue, 2020). Here our data show that if the liver is inflamed prior to SCI, the harmful consequences of SCI on the liver are exacerbated. That is, rBDL alone caused transient hepatic inflammation, and SCI alone caused a persistent rise in serum ALT, acute phase response proteins (APPs) and hepatic cytokines. The combination of rBDL+SCI, however, potentiated liver damage, as noted by enhanced serum ALT, hepatic cytokine expression, Kupffer cell activation, iron and lipid accumulation, and hepatocyte ballooning.

Fetuin-A is another feasible mediator of SCI-induced NASH and liver pathology. Fetuin-A production can be induced by NFkB activation (Dasgupta *et al*., 2010), which is stimulated by factors elevated in the liver after SCI, including TNFα. Fetuin-A can stimulate SREBP-1c expression as well as mediate palmitate-induced steatosis (Jung *et al*., 2013); thus, it may have contributed to the enhanced hepatic lipid accumulation detected in this study. As stated above, Fetuin-A acts as an endogenous TLR4 ligand to promote pro-inflammatory signaling; it also inhibits insulin receptor signaling (Pal *et al*., 2012; Jung, Yoo and Choi, 2016; Lebensztejn *et al*., 2016). Indeed, Fetuin-A is thought to be a possible link between NAFLD, obesity and insulin resistance (Lebensztejn *et al*., 2016). Since these are common clinical features of SCI (Bauman and Spungen, 2001; Manns, McCubbin and Williams, 2005; LaVela *et al*., 2006), Fetuin-A may be an important mediator of hepatic pathology as well as development of metabolic syndrome after SCI.

Another possible mechanism driving hepatic pathology after SCI is overactive sympathetic nervous system (SNS) signaling to the liver. The mid-thoracic injury model used here disrupts most of the descending inhibitory control over intraspinal sympathetic neurons that regulate liver function. This “disinhibition” leads to excess systemic SNS firing after SCI, which in the liver causes Kupffer cell activation and increased inflammatory cytokine expression, including a 4-7-fold increase in TNFα (Zhou *et al*., 2001; Liu *et al*., 2012; Lin *et al*., 2015). Within the liver, TNFα can promote lipid accumulation by increasing expression of lipid uptake transporters such as CD36 and lipogenic molecules like serine palmitoyltransferase (SPT) and sterol regulatory response element bind protein-1c (SREBP-1c) (Endo *et al*., 2007; Pagadala *et al*., 2012; Martius *et al*., 2015). Accordingly, a recent study using high fat diet-induced NAFLD showed inhibiting TNF receptor-1 reduced steatosis, SREBP-1c expression, overall indices of liver pathology (Wandrer *et al*., 2020). Therefore, after SCI, SNS-induced TNFα expression may be a culprit in the elevated liver pathology.

Hepatic inflammation after SCI could also have been induced by bacterial translocation from the gastrointestinal (GI) tract to the liver. Prior work from our group showed SCI causes rapid and robust gut dysbiosis and bacterial translocation in mice (Kigerl *et al*., 2016; Kigerl, Mostacada and Popovich, 2018). Here we confirm circulating endotoxin levels rise after SCI and show that they are even higher in rats receiving SCI concomitant with liver inflammation. Transient bile duct ligation may increase intestinal permeability and bacterial translocation (Fouts *et al*., 2012), and here, rBDL alone increased circulating endotoxins starting 5 days after ligation reversal which declined by the end of the study. However, the combination of SCI plus rBDL clearly potentiates the response to both. Multiple hepatic parenchymal cells express TLR4, including hepatocytes, Kupffer cells and stellate cells. Thus, prolonged exposure to gut-derived bacterial products after SCI, as occurred in this study, could feasibly induce and sustain hepatic inflammation.

Lastly, other potential though less likely mediators of enhanced liver (or spinal) pathology are serum bile salts and ammonia, which were elevated for 5 days after ligation reversal in rBDL+SCI rats. Previous work showed that even transient elevation of circulating ammonia or bile can affect liver steatosis and inflammation in humans and rodents (Lake *et al*., 2013; Tranah *et al*., 2013; Ferslew *et al*., 2015; Puri *et al*., 2018). In our study rBDL alone significantly increased liver fat droplet size and caused a 2-fold increase Kupffer cell activation at 23dpi compared to controls (although not at high as in the rBDL+SCI group). This is mirrored by studies showing that NAFLD/NASH are positivity correlated with increased bile acid synthesis (Boursier and Diehl, 2015; Ferslew *et al*., 2015; Mouzaki *et al*., 2016).

### Clinical relevance of hepatic inflammation at the time of SCI

Clearly most people sustaining SCI do not present with concomitant bile duct blockade or cholecystitis. Here reversible bile duct ligation was used as a tool to induce hepatic inflammation reliably and consistently. However, a large percentage of people likely do have inflamed livers at the time they sustain SCI. For instance, a sizeable portion of the population is overweight or obese and, accordingly, a clinical study reported that at least 25% of acute SCI patients are obese; notably, these individuals had significantly worse outcomes compared to non-obese SCI victims (Stenson *et al*., 2011). Obesity is widely recognized to be associated with hepatic inflammation and lipid accumulation. Indeed, NASH is considered the hepatic manifestation of metabolic syndrom (Farrell and Larter, 2006; Dietrich and Hellerbrand, 2014). Thus, obese patients sustaining SCI will likely be injured on a background of liver inflammation similar to the rats studied here. Another obvious condition associated with liver damage is alcoholism. Notably, alcohol is a factor in a large proportion of SCIs (Heinemann *et al*., 1988; Kiwerski and Krasuski, 1992) and individuals with history of alcohol abuse have poorer outcomes from injury (Hawkins and Heinemann, 1998; Elliott *et al*., 2002). SCI in this population again would occur on a background of hepatic inflammation and pathology. A third important group to consider is the aging population. Age is associated with increased liver inflammation (Williams *et al*., 2011; Bertolotti *et al*., 2014) and statistics show that the number of elderly individuals sustaining SCI has been rising for several years (Devivo and Chen, 2011; Devivo, 2012; Chen, He and DeVivo, 2016). While this group is expected to have diverse underlying conditions at the time of injury, hepatic inflammation is likely in at least a subset of elderly patients. Thus, it is critical to understand how pre-existing liver inflammation due to any number of conditions results in worse hepatic and intraspinal damage and reduced functional recovery after SCI. This is especially true in light the current data showing the devastating effects of combining permanent bile duct ligation with SCI, which suggests patients presenting with significant liver pathology at the time of injury may be particularly vulnerable.

### Potential metabolic dysfunction related to liver pathology after SCI

Individuals with SCI commonly develop features of metabolic syndrome, including increased adiposity, insulin resistance and hyperlipidemia, which collectively increase their risk for heart disease and diabetes (Kocina, 1997; Bauman and Spungen, 2001, 2008; Spungen *et al*., 2003; LaVela *et al*., 2006; Weaver *et al*., 2007; Inskip *et al*., 2010; Rankin *et al*., 2017). Evidence suggests that NASH contributes to these conditions and is an important factor in the increased risks of disease after SCI (Ballestri *et al*., 2016; Hazlehurst *et al*., 2016; Lonardo *et al*., 2016; Mahfood Haddad *et al*., 2017). For example, 50% of individuals with metabolic syndrome have NAFLD, and liver fat content is strongly correlated with the number of metabolic syndrome features present (Kotronen *et al*., 2007; Lonardo *et al*., 2015). Specifically, hepatic steatosis is a strong predictor of insulin resistance in skeletal and cardiac muscle and adipose tissue (Agosti, Sabbà and Mazzocca, 2018). Indeed, ∼50% of chronic SCI subjects display NAFLD along with insulin resistance (Sipski *et al*., 2004; Shin *et al*., 2006; Barbonetti *et al*., 2016). Taken together, these previous studies and our current results suggest increased hepatic inflammation and steatosis may be a link to the increased metabolic dysfunction, diabetes and cardiovascular disease observed in the SCI population.

## Conclusions

The mechanisms driving liver pathology and metabolic dysfunction after SCI remain largely elusive and poorly understood. The current work provides the first evidence of SCI-induced perturbation of a liver-spinal cord axis in a bi-directional manner that drives chronic pathology in both organs, enhances SCI-induced metabolic dysfunction and reduces functional recovery. These studies suggest a novel potential role for Fetuin-A in driving this disruption and suggest that the liver and its products could be feasible clinical targets for reducing symptoms of hepatic metabolic dysfunction and improving overall recovery from SCI.

## Acknowledgements

The authors thank Feng Qin Yin, Rochelle Deibert, Ping Wei, Zhen Guan and Yaoling Shu for excellent technical assistance. We also thank the DHSB for supplying the anti-neurofilament antibody. This work was funded by the National Institute of Neurological Disorders and Stroke (NINDS) P30-NS045758 (D.M.M.), R01-NS082095 (D.M.M.), Craig H. Neilsen Foundation (M.G.) and the Belford Center for Spinal Cord Injury.

## Author Contributions

M.T.G., A.D.S., and D.M.M designed and performed the experiments and prepared the figures; M.T.G., K.E.C., A.D.S., P.D., and A.D.A contributed to the performance of the experiments; All authors assisted in preparing the manuscript; and M.T.G., R.S.B., P.G.P. and D.M.M. supervised the work and wrote the manuscript.

